# Quantitative mass imaging of single molecules in solution

**DOI:** 10.1101/229740

**Authors:** Gavin Young, Nikolas Hundt, Daniel Cole, Adam Fineberg, Joanna Andrecka, Andrew Tyler, Anna Olerinyova, Ayla Ansari, Erik G. Marklund, Miranda P. Collier, Shane A. Chandler, Olga Tkachenko, Joel Allen, Max Crispin, Neil Billington, Yasuharu Takagi, James R. Sellers, Cedric Eichmann, Philip Selenko, Lukas Frey, Roland Riek, Martin R. Galpin, Weston B. Struwe, Justin L.P. Benesch, Philipp Kukura

## Abstract

The cellular processes underpinning life are orchestrated by proteins and their interactions. Structural and dynamic heterogeneity, despite being key to protein and drug function, continues to pose a fundamental challenge to existing analytical and structural methodologies used to study these associations. Here, we use interferometric scattering microscopy to mass-image single biomolecules in solution with <2% mass error, up to 19-kDa resolution and 1-kDa precision. Thereby, we resolve oligomeric distributions at high dynamic range, detect small-molecule binding, and mass-image biomolecules composed not only of amino acids, but also heterogeneous species, such as lipo- and glycoproteins. These capabilities enable us to characterize the molecular mechanisms of processes as diverse as oligomeric self-assembly, glycoprotein cross-linking, amyloidogenic protein aggregation, and actin polymerization. Interferometric scattering mass spectrometry (iSCAMS) provides spatially resolved access to the dynamics of biomolecular interactions ranging from those involving small molecules to mesoscopic assemblies, one molecule at a time.

---

Macromolecular complexes perform diverse tasks inside the cell, such as the translation of the genetic code (*1*), energy conversion (*2*), transport (*3*), cell migration and communication (*4*). These cellular machines almost exclusively consist of a few to several thousand individual components, including proteins, nucleic acids, carbohydrates and lipids, held in shape by non-covalent interactions (*5*). In most cases, proteins only become active as a consequence of association with binding partners. In others, assembly and disassembly, be it native or regulated through external factors, results in the desired activity (*6*).

Attempts to study the mechanisms underlying protein function have thus largely focused on two approaches: the development of ever more powerful structural techniques with atomic resolution (*7*) and the application of advanced ensemble methodologies aimed at unraveling assembly and disassembly (*8*). At the single molecule level, significant progress has been made in terms of characterizing interactions (*9*) and dynamics (*10*, *11*), but there remains a lack of universally applicable techniques capable of addressing the range of interactions involving few to thousands of molecules.

Given sufficient sensitivity, light scattering is an ideal candidate for detecting and characterizing biomolecules, because of its universality. For an interferometric detection scheme (Fig. 1A), the scattering signal scales with the polarizability, which is a function of the refractive index and proportional to the particle volume (*12*). Combining the approximation that single amino acids effectively behave like individual nano-objects with the observation that the specific volumes of amino acids and refractive indices of proteins vary by only ~1% (Supplementary Fig. 1; Supplementary Table I) suggests that the number of amino acids in a polypeptide and thus mass is proportional to scattering signal. This close relationship between mass and interferometric contrast, which has been predicted (*13*, *14*) and coarsely observed (*15*, *16*) to hold even at the single molecule level, could thus in principle achieve a mass accuracy comparable with native mass spectrometry given sufficient detection sensitivity.

**Fig. 1.**
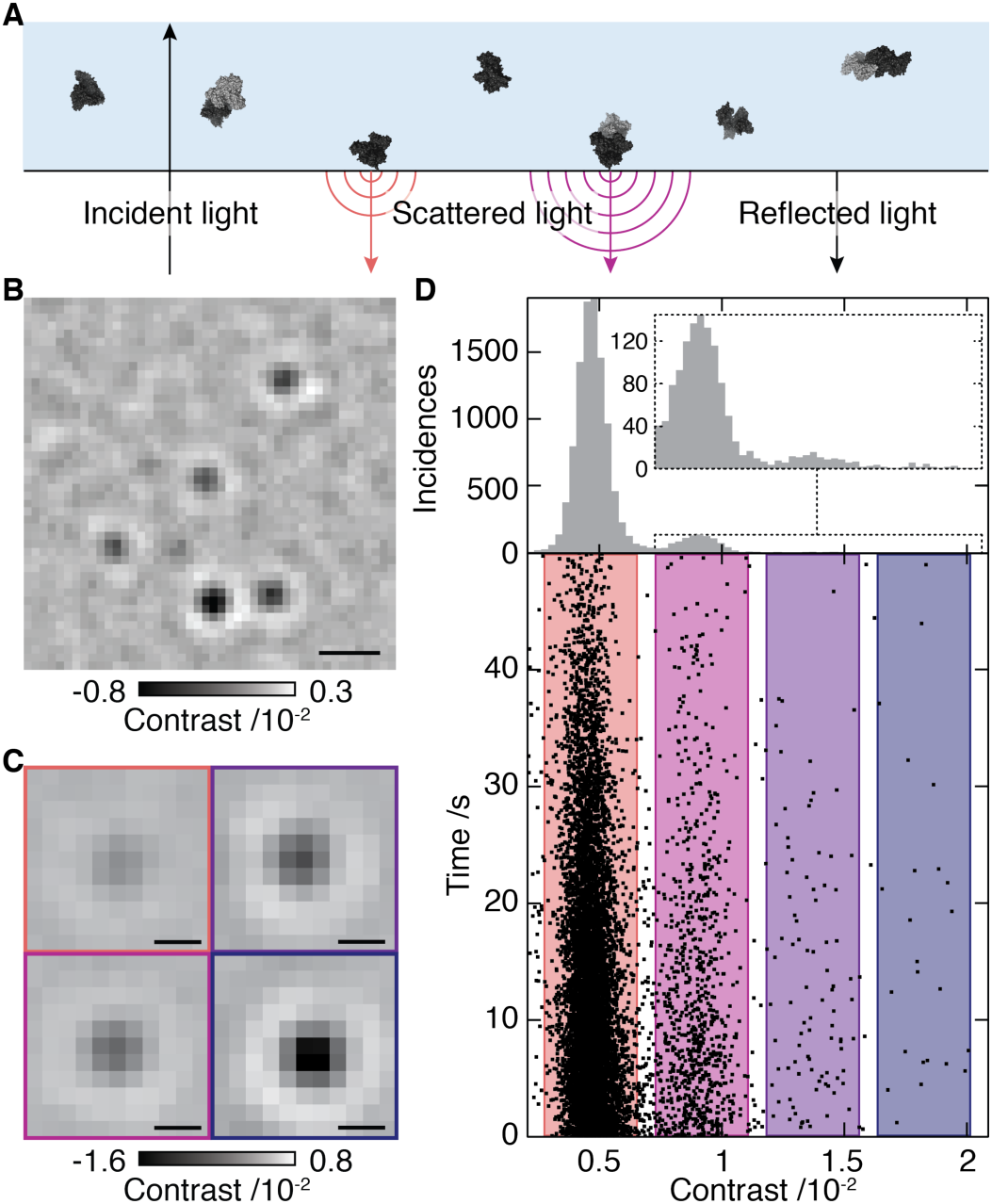
Concept of interferometric scattering mass spectrometry (iSCAMS). **(A)** Schematic of the experimental approach. **(B)** Differential interferometric scattering image of BSA. Scale bar: 0.5 μm. **(C)** Representative images of monomers, dimers, trimers and tetramers of BSA. Scale bar: 200 nm. **(D)** Scatter plot of single molecule binding events and their scattering contrasts for 12 nM BSA from 14 movies (lower). Corresponding histogram (N=11482), and zoom of the region for larger species (upper). The reduction in landing rate results from a drop in BSA concentration with time.

Building on recent advances in signal detection that enable higher imaging contrast for interferometric scattering microscopy (*16*, *17*), we could obtain high quality images of single proteins binding non-specifically to a microscope coverslip (Fig. 1B, Supplementary Movie 1). Reaching signal-to-noise ratios >10, even for small proteins such as bovine serum albumin (BSA), combined with an optimized data analysis approach (*17*) allowed us to accurately extract the scattering contrast for each molecular binding event (Fig. 1C, Supplementary Fig. 2). These improvements enabled us to reveal clear signatures of different oligomeric states, here shown for BSA with relative abundances of 88.63%, 9.94%, 1.18% and 0.25% (Fig. 1D). These results demonstrate the ability of iSCAMS to characterize oligomeric distributions of biomolecules in solution and high dynamic range with respect to the detection and quantification of rare complexes, as evidenced by observation of BSA tetramers.

The regular spacing in the contrast histogram of BSA tentatively confirms the expected linear scaling between mass and interferometric contrast. Repeating these measurements for eight different proteins, spanning 53 – 803 kDa, revealed a linear relationship with an average error of 1.9% (Fig. 2A, Supplementary Fig. 3A). The resolution, as defined by the full-width at half-maximum (FWHM) of the measured contrast reached 19 kDa for streptavidin. In all cases, the resolution was not only limited by shot noise (Supplementary Fig. 4A), but also influenced by molecular mass, increasing from 19 kDa for streptavidin to 102 kDa for thyroglobulin (Supplementary Fig. 4B), due to local variations in coverslip roughness. Deviations between measured and sequence mass could not be attributed to overall molecular shape (Supplementary Fig. 4C), and may instead be caused by factors such as the solvation environment and binding mode, which become relatively less important for larger proteins. Even for large structural differences, such as those between the extended and folded conformation of smooth-muscle myosin (530.6 kDa, Fig. 2A and Supplementary Fig. 3B,F), we did not find measurable differences in the molecular mass beyond the mass increase expected for addition of many glutaraldehyde molecules (Extended conformation: 528.4 ± 16.2 kDa, folded conformation: 579.4 ± 14.8 kDa), which were used to crosslink and thus lock myosin in the folded conformation. The sub-0.5% deviation from sequence mass for species of >100 kDa compares well to native mass spectrometry (*18*), and demonstrates the intrinsic capability of iSCAMS as an accurate mass measurement tool for biomolecules.

**Fig. 2.**
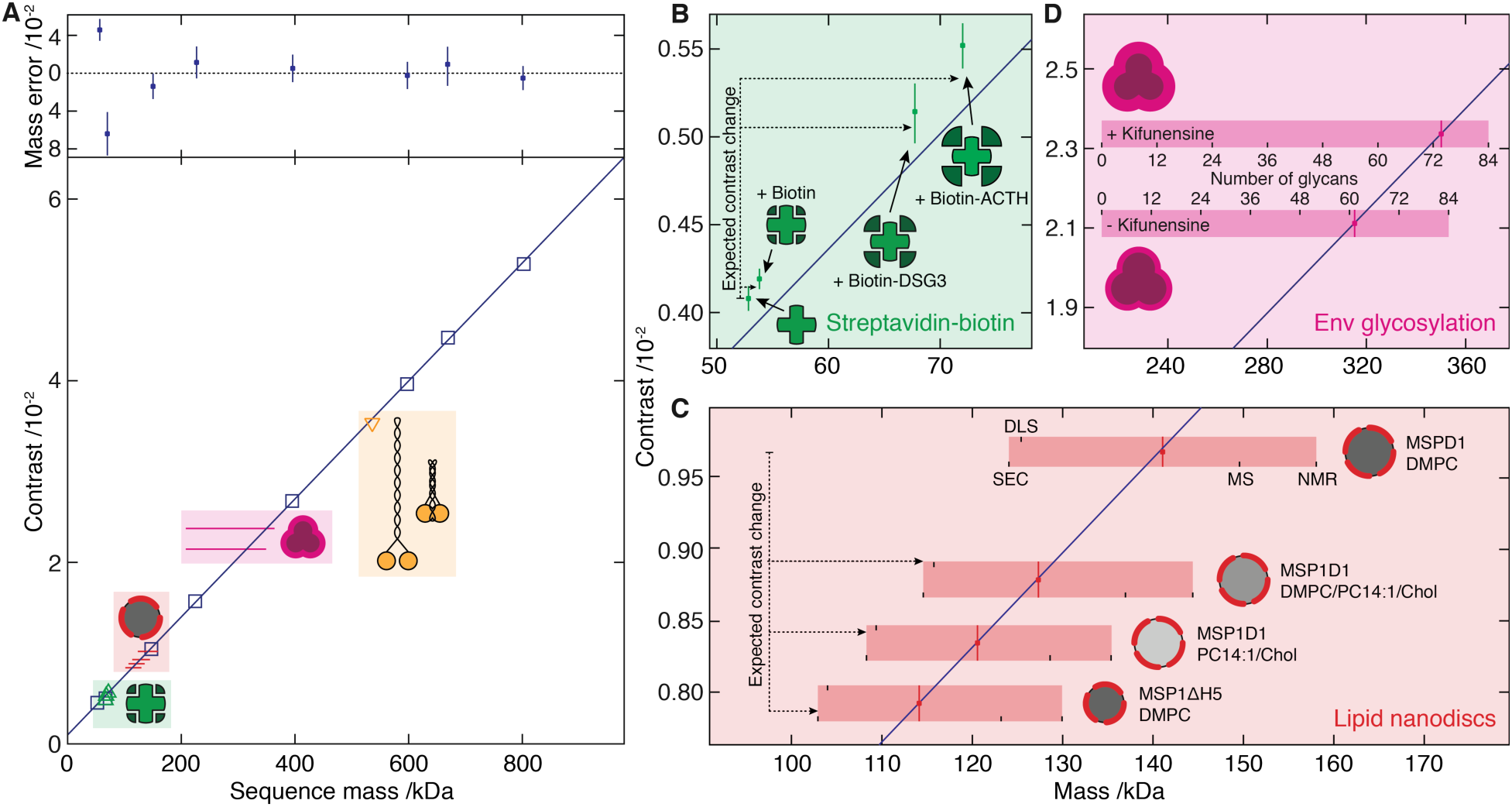
Characterization of iSCAMS accuracy, precision, and dependence on molecular shape and identity. **(A)** Contrast vs molecular mass including proteins used for mass calibration (blue), and characterization of shape dependence (yellow), protein-ligand binding (green), lipid nanodisc composition (red) and glycosylation (purple). Mass error (upper panel) is given as a percentage of the sequence mass relative to the given linear fit. **(B)** Mass-sensitive detection of ligand binding using the biotin-streptavidin system. **(C)** Nanodisc mass-measurement for different lipid compositions and protein belts. Masses obtained by alternative methodologies for MSP1D1/DMPC are marked, and extrapolated to the other compositions. **(D)** Recorded differential contrast for Env expressed in the presence or absence of kifunensine, and associated mass ranges expected for different glycosylation levels. Error bars represent the standard error of the mean from repeated experiments.

The high precision of 1.8 ± 0.5% with respect to the protein mass (Fig. 2A), likely also limited by coverslip roughness, indicates the potential for direct detection of small-molecule binding. We therefore examined the biotin-streptavidin system (Fig. 2B, Supplementary Fig. 3C), and measured masses for streptavidin in the absence (55.7 ± 1.1 kDa) and presence (57.4 ± 0.9 kDa) of a 100-fold excess of biotin. This corresponds to a difference of 1.7 ±1.4 kDa, in good agreement with the expected 0.98 kDa for complete occupancy of the four binding sites. Upon addition of two different biotinylated peptides (3705.9 Da and 4767.4 Da), we obtained increases of 16.1 ± 2.8 kDa and 22.0 ± 2.2 kDa (compared to 14.8 kDa and 19.1 kDa expected) (Fig. 2B, Supplementary Fig. 3C). These data demonstrate that iSCAMS can detect the association of kDa-sized ligands, making it suitable for highly sensitive, mass-sensitive ligand-binding studies in solution.

The results for biotin-binding led us to question to what degree iSCAMS can be extended to characterize biomolecules containing components other than amino acids. Lipid nanodiscs, high-density engineered lipoprotein particles, represent an ideal test system due to their flexibility in terms of polypeptide and lipid content (*19*). For nanodiscs composed of the MSP1D1 belt protein and DMPC lipids, we obtained a mass of 141.0 ± 1.6 kDa, in good agreement with the range of masses reported by other methods, spanning 124 – 158 kDa (Fig. 2C and Supplementary Fig. 3D). Replacing MSP1D1 with the smaller MSP1DH5 reduces the nanodisc diameter and the lipid content by ~20%, after accounting for the thickness of the protein belt (*20*). Given the masses of MSP1D1 and MSP1DH5 (47 and 42 kDa, respectively), we predicted a mass for the MSP1DH5 nanodisc of 113.6 kDa, in excellent agreement with our measurement (114.1 ± 1.9 kDa). Notably, mass shifts associated with changes in lipid composition, such as those introduced by partially unsaturated lipids and cholesterol, matched those predicted from the assembly ratios (Fig. 2C, Supplementary Tables II-VI).

To see whether our approach also applies to solvent-exposed chemical species that experience a different dielectric environment to those buried within a protein, we selected the HIV envelope glycoprotein complex (Env), which is a trimer of gp41-gp120 heterodimers. Env is extensively N-glycosylated, with the carbohydrates contributing to almost half of its mass (*21*). For an Env trimer mimic expressed in the presence of kifunensine, a mannosidase inhibitor that leads predominantly to unprocessed Man_9_GlcNAc_2_ glycans (Supplementary Fig. 5), we recorded a mass of 350.0 ± 5.7 kDa. Making the crude approximation that glycans and amino-acids have similar polarizabilities, this corresponds to a glycan occupancy of 74 ± 3 out of 84 possible sites (Fig. 2D and Supplementary Fig. 3E), consistent with recent observations of high occupancy for gp120 expressed with kifunensine (*22*). For Env expressed without kifunensine we recorded a lower mass of 315.3 ± 10.5 kDa. The mass difference can only in part be attributed to the lower average mass of the processed glycans (Supplementary Fig. 5), and yields a total N-glycan occupancy of 61 ± 6. While the exact values for occupancy are beholden to our calibration (Fig. 2A), the presence of unoccupied sites is consistent with their observation in proteomics data (*23*). Taken together, these experiments demonstrate that iSCAMS is applicable to the mass measurement of different classes of biomolecules, including assemblies comprising heterogeneous mixtures thereof, which are particularly challenging to existing analytical techniques.

Monitoring protein assembly and detecting all the involved species in order to reveal the underlying molecular mechanisms and energetics is a significant challenge, but one that iSCAMS is well suited for, especially at the low concentrations often required to reach disassembly under native conditions. To illustrate this, we chose HSP16.5 from the hyperthermophile *Methanocaldococcus jannaschii* (*24*), which forms a highly stable 24-mer assembled from 12 dimers. Upon decreasing the HSP16.5 concentration, we observed clear signatures of decomposition into smaller oligomers, with an increasing abundance of small compared to large species (Fig. 3A). A minimal model assuming identical equilibrium constants for all oligomers except for the 24-mer, with *K′_Dimer_* = 130 *K_Dimer_,* reproduced the experimental oligomeric evolution remarkably well, suggesting that HSP16.5 assembles without any stable intermediates, and the final product being energetically preferred by ~5 kJ mol^−1^.

**Fig. 3.**
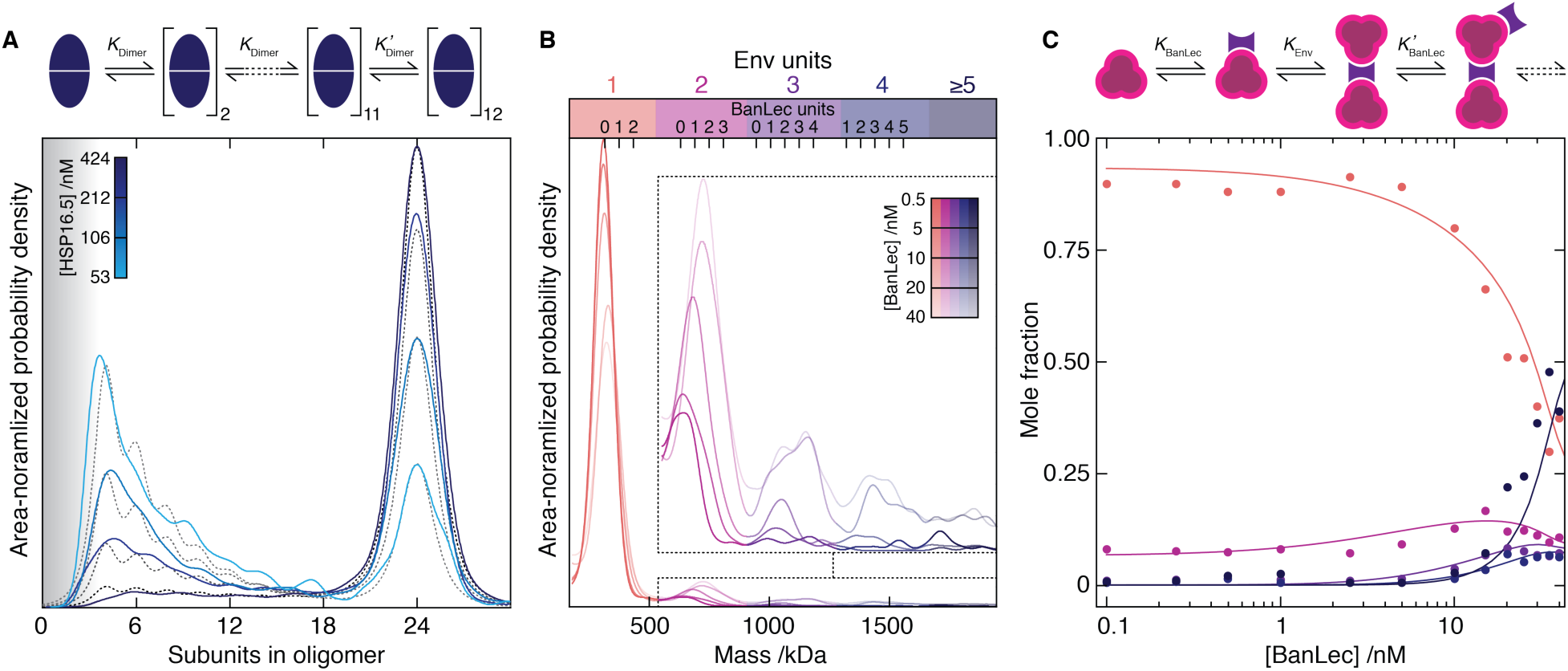
Single molecule mass spectrometry of protein aggregation and disassembly. **(A)** Evolution of the mass distribution as a function of HSP16.5 concentration (solid) and corresponding fit (dashed) to a consecutive assembly mechanism assuming identical equilibrium constants for all assembly intermediates except for the last, where *K′_Dimer_* = 130 × *K_Dimer_.* The grey shading indicates a region of low detection sensitivity. **(B)** Mass distributions for Env in the presence of 0.5 − 40 nM BanLec monomer. Inset: zoom alongside expected positions for multiples of bound BanLec tetramers. **(C)** Oligomeric fractions colored according to **B** vs BanLec concentration including predictions (solid) using the given cooperative model.

To test whether iSCAMS is capable of characterizing more complex assembly processes, we investigated the interaction of Env with the anti-viral lectin BanLec, which is known to neutralise HIV by binding to surface N-glycans (*25*) via an unknown mechanism. Addition of BanLec to Env resulted in a reduction of single Env units coupled to the appearance of dimers and higher-order assemblies (Fig. 3B). Close inspection of the mass distributions revealed shifts consistent with stoichiometric binding of BanLec to Env, as opposed to coating of the Env surface with BanLec (Fig. 3B, inset). A model based on cooperative binding of BanLec matched the recorded oligomeric evolution closely (Fig. 3C), allowing us to extract the respective association constants (*K_Env_* = 8 nM^−1^, K′_BanLec_ = 0.4 nM^−1^, K_BanLec_ = 0.12 nM^−1^). Using iSCAMS we were therefore able to extract the energetics underlying the lectin-glycoprotein interaction, despite the heterogeneity of this multi-component system, and anticipate similar quantitative insights to be achievable for other therapeutic target proteins.

An advantage of our imaging-based approach stems from its ability to time-resolve mass changes in a position- and local concentration-sensitive manner. This enables us to examine surface-catalyzed nucleation events, for example those involved in amyloid formation (*26*). Upon adding the amyloidogenic protein α-synuclein to a planar, negatively charged DOPC/DOPS (3:1) bilayer at physiological pH, we observed the appearance and growth of nanoscopic objects within seconds even at low μM concentrations (Fig. 4A, Supplementary Movie 2). Growth of these nucleation sites was uniform across the field of view, with the initial rates following approximately first-order kinetics (Fig. 4B and Supplementary Fig. 6A), pointing towards a simple growth mechanism. We did not detect such structures on neutral, DOPC-only bilayers, and found evidence for thioflavin-T positive aggregates after overnight incubation (Supplementary Fig. 6B), suggesting that our assay probes the early stages of amyloid assembly.

**Fig. 4.**
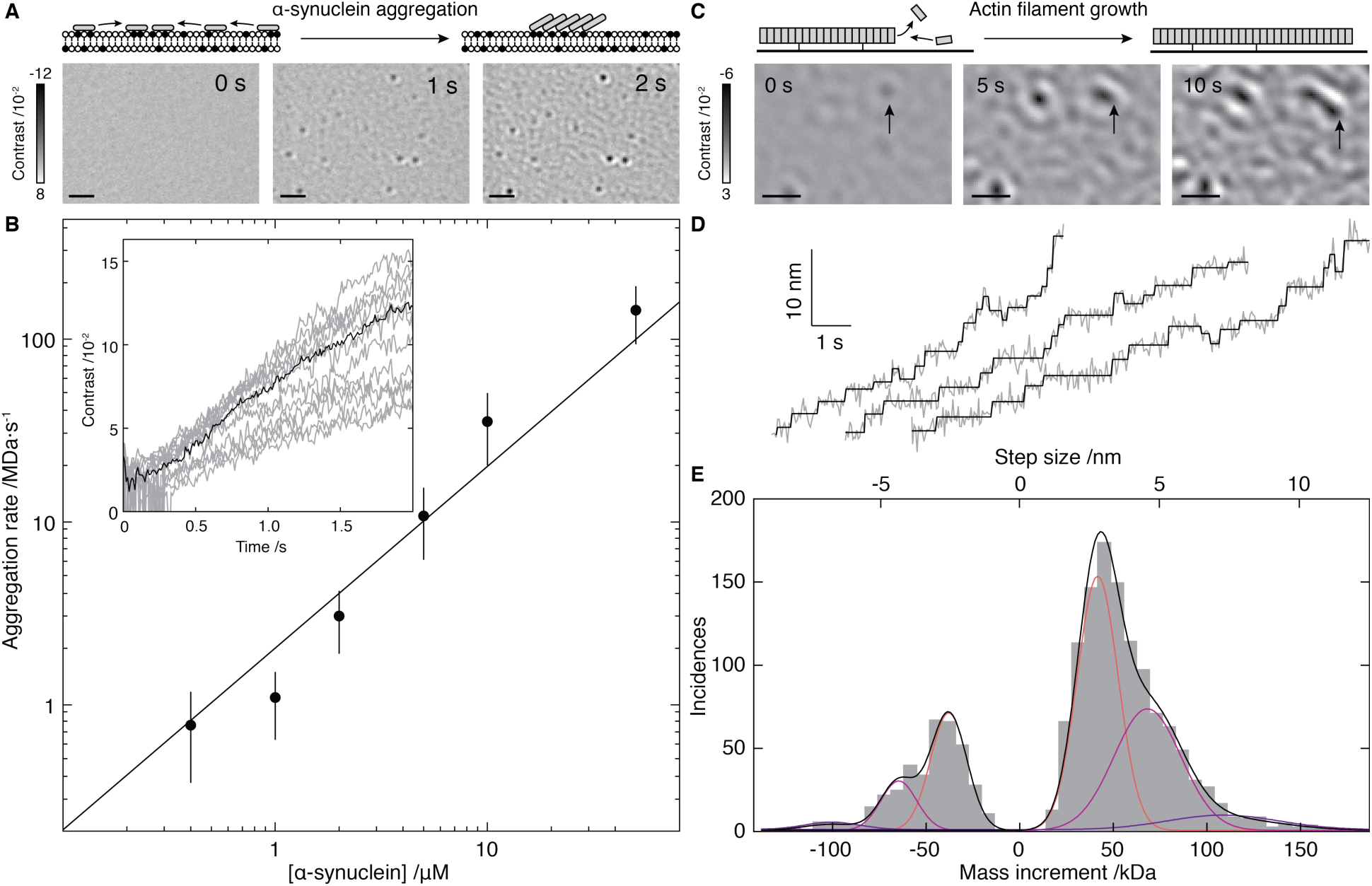
Mass-imaging of mesoscopic dynamics. **(A)** Schematic of and iSCAMS images for α-synuclein (1 μM) aggregation on a negatively charged bilayer membrane. **(B)** Initial growth rate vs. α-synuclein concentration alongside expectations from first order kinetics (solid). Inset: Individual growth trajectories (grey) and average (black) for 21 particles from **A**. **(C)** Schematic and iSCAMS images of actin polymerization. The arrow highlights a growing filament. **(D)** Representative traces of actin filament tip position (grey) and corresponding detected steps (black). **(E)** Step and mass histogram from 1523 steps and 33 filaments including a fit to a Gaussian mixture model (black) and individual contributions (colored). Scale bars: 1 μm.

At the extremes of our current sensitivity, iSCAMS enables mass-imaging of mesoscopic self-assembly, molecule-by-molecule. In an actin polymerization assay, subtraction of the constant background revealed growth from surface-immobilized filaments, and their length changes upon the attachment and detachment of actin subunits (Fig. 4C, Supplementary Fig. 7A, Supplementary Movie 3). We observed distinct changes in the filament length (Fig. 4D, Supplementary Fig. 7B-D and Supplementary Movie 4), the most frequent forward and backward step sizes in the traces being 3.0 ± 0.8 nm and 2.7 ± 0.7 nm, respectively, remarkably close to the expected length increase of 2.7 nm upon binding of a single actin subunit to a filament (Fig. 4E). Detection of larger step sizes represents the addition of multiple actin subunits within our detection time window. The contrast changes associated with the different step sizes corresponded to mass changes of one, two, or three actin monomers binding and unbinding to the tip of the growing filaments during acquisition (Supplementary Fig. 7E,F). Even though we cannot yet distinguish between models invoking monomer (*27*) or oligomer (*28*) addition to a growing filament at our current level of spatio-temporal resolution, these results demonstrate the unique capability of iSCAMS for quantitatively imaging biomolecular assembly at the single molecule level.

We envision that the combination of experimental simplicity and broad applicability of iSCAMS will make it a powerful approach for identifying biomolecules and quantifying their interactions. The technology will enable insights into self-assembly at the single molecule level, and the determination of associated thermodynamic and kinetic parameters. Combining iSCAMS with recent developments that are compatible with optical microscopy, such as passive trapping of single molecules (*29*) and microfluidics (*30*), will provide access to further improvements in sensitivity, accuracy, resolution and scope of single molecule mass measurement in solution.

## Acknowledgments

GY was supported by a Zvi and Ofra Meitar Magdalen Graduate Scholarship; NH by a DFG research fellowship (HU 2462/1-1). EGM thanks the Swedish Research Council and the European Commission for a Marie Skłodowska Curie International Career Grant (2015-00559). MPC is a Clarendon Scholar supported by the Oxford University Press. SAC is supported by the Biotechnology and Biological Sciences Research Council and Waters Corp by the iCASE studentship BB/L017067/1 to JLPB, JA and MC were supported by the National Institute of Allergy and Infectious Diseases (Center for HIV/AIDS Vaccine Immunology and Immunogen Discovery grant UM1AI100663). JRS is supported by NHLBI Intramural program HL0001786 and thanks Fang Zhang for technical assistance and the NHLBI electron microscopy core facility. CE is supported by a SNSF advanced postdoctoral mobility fellowship (P300PA160979). PS is funded by an ERC Consolidator Grant (NeuroInCellNMR 647474). JLPB thanks the Engineering and Physical Sciences Research Council for EP/J01835X/1. PK was supported by an ERC Starting Investigator Grant (Nanoscope, 337577).

Conceptualization, WBS, JLPB and PK; Methodology GY, NH, DC, JA, EGM, CE, PS, MRG, WBS, JLPB, PK; Software GY, NH; Validation GY, NH, JLPB, PK; Formal analysis, GY, NH, AT, AA, AO, JA, EGM, MRG; Investigation GY, NH, DC, AF, JA, AT, AA, NB, YT, CE; Resources MPC, SAC, OT, JA, MC, NB, YT, JRS, CE, PS, LF, RR, WS; Writing - original draft GY, JLPB, PK; Writing - review and editing GY, NH, AF, AO, EGM, MPC, OT, MC, JRS, CE, PS, RR, MRG, WBS, JLPB, PK; Visualization GY, NH, JLPB, PK; Supervision PK.

## Supplementary Materials

Materials and Methods

Figures S1-S7

Tables SI-SVI

Movies S1-S4 References (*31-52*)

